# Dose-dependent phorbol 12-myristate-13-acetate-mediated monocyte-to-macrophage differentiation induces unique proteomic signatures in THP-1 cells

**DOI:** 10.1101/2020.02.27.968016

**Authors:** Sneha M. Pinto, Hera Kim, Yashwanth Subbannayya, Miriam Giambelluca, Korbinian Bösl, Richard K. Kandasamy

**Author notes:** Correspondence: Richard K. Kandasamy. These authors contributed equally (Sneha M. Pinto, Hera Kim).

## Abstract

Macrophages are sentinels of the innate immune system, and the human monocytic cell line THP-1 is one of the widely used *in vitro* models to study immune responses. Several monocyte-to-macrophage differentiation protocols exist, with phorbol 12-myristate-13-acetate (PMA) being the widely used and accepted method. However, the concentrations and duration of PMA treatment vary widely in the published literature and their effect on protein expression is not fully deciphered. In this study, we employed a dimethyl labeling-based quantitative proteomics approach to determine the changes in the protein repertoire of macrophage-like cells differentiated from THP-1 monocytes by three commonly used PMA-based differentiation protocols. Our analysis shows that variations in PMA concentration and duration of rest post-stimulation result in downstream differences in the protein expression and cellular processes. We demonstrate that these differences result in altered gene expression of cytokines upon stimulation with various TLR agonists. Together, these findings provide a valuable resource that significantly expands the knowledge of protein expression dynamics with one of the most common *in vitro* models for macrophages, which in turn has a profound impact on the immune responses being studied.

## Introduction

Macrophages and their precursors- monocytes, mediate innate immune responses and inflammatory processes, and also contribute to adaptive immunity through antigen presentation (1, 2). Monocytes in the blood circulation migrate to the site of infection/inflammation and differentiate into macrophages for effective host defense, tissue remodeling, and repair (3). Furthermore, macrophages exhibit a high level of plasticity, depending on their local microenvironment, specialized functions and varied phenotype acquired (1, 4).

Several models are employed to study the mechanisms of immune-modulation in monocytes and macrophages during inflammatory diseases. The most frequently used include primary peripheral blood mononuclear cells (PBMCs) and monocyte cell lines. However, due to donor-to-donor variations and technical disparities involved in handling of PBMCs *in vitro*, the human leukemia monocytic cell line, THP-1, is widely accepted and used as a monocyte/macrophage model (5, 6). Several studies have indicated that THP-1 cells can be differentiated into macrophage-like cells using phorbol-12-myristate-13-acetate (PMA), which markedly resembles PBMC monocyte-derived macrophages (MDMs) in cytokine production, metabolic and morphological properties, including differential expression of macrophage surface markers such as *CD14*, CD11b (*ITGAM*) and scavenger receptors- *CD163, MSR1*, and *SCARB2* (5, 7-10). Nonetheless, depending on the parameters of the differentiation protocol employed, such as the concentration (ranging from 5 to 400 ng/mL), and duration of incubation (1 to 5 days) with PMA; the degree of differentiation and the functional changes may vary significantly (5, 11-15).

At the molecular level, multiple proteins including growth factors, antigenic markers, chemokine-receptors, cytokines, and cell adhesion molecules, are known to govern and reflect underlying monocyte-macrophage differentiation processes (16-20). However, the effect of various differentiation protocols on the cellular proteome and intracellular signaling networks during monocyte-to-macrophage differentiation remains poorly understood. Hence, it is crucial to determine the most suitable differentiation conditions when using these cells as model system, as this can significantly impact their response to various innate immune stimuli. Quantitative high-resolution mass spectrometry-based proteomic approaches have been widely employed to investigate the proteomes of monocytes and macrophages as well as altered cellular proteomes and complex cellular/biological mechanisms in several biological conditions (21-23). However, to date, no studies have directly compared the proteome changes of the PMA-mediated differentiation process.

In the present study, we evaluate the effect of three PMA-based differentiation protocols on the changes in the proteome profiles upon THP-1 differentiation using stable isotope dimethyl labeling quantitative proteomics. We demonstrate that various differentiation conditions such as concentration and incubation time, prior to any stimuli, are critical consideration factors for heterogeneity of the cell culture. The results from this study will enable immunologists to make informed decisions on differentiation protocols that result in the desired proteotypes for custom experiments.

## Material and methods

### Cell culture and differentiation

Human THP-1 monocytic cells (ATCC) were cultured in RPMI 1640 (Sigma-Aldrich) medium containing 10% heat-activated fetal calf serum (FCS), 2 mM L-glutamine, 100 nM penicillin/streptomycin (Thermo Fisher Scientific) and 50 μM *β*-mercaptoethanol (Sigma-Aldrich). The cells were maintained in a humidified 37°C, 5% CO_2_ incubator. THP-1 cells were differentiated into resting macrophages by resuspending the cells in growth medium containing 5 or 50 ng/ml phorbol-12-myristate-13-acetate (PMA; Sigma-Aldrich) and cultured for indicated time periods. The process of differentiation was enhanced by removing the PMA-containing media and adding fresh, complete RPMI 1640 media to the cells. The treatment conditions included the following. **Condition A**: 50 ng/ml PMA for 72 hours followed by 48 hours rest (5 days); **Condition B**: 50 ng/ml PMA overnight (16 hours) followed by 48 hours rest (Condition B); and **Condition C**: 5 ng/ml PMA for 48 hours followed by 3 hours rest (5, 11-14).

### Phase-contrast microscopy

THP-1 cells were seeded at a density of 0.2 × 10^6^ cells/ml in cell culture Cellvis glass-bottom plates and treated with PMA and rested at indicated concentrations and duration. The morphological characteristics of undifferentiated and differentiated THP-1 cells were captured by EVOS FL Auto Cell Imaging System 2 (Thermo Fisher Scientific) using a 40x objective lens and were processed by ImageJ software (W.S. Rasband, National Institutes of Health, Bethesda, MD).

### Flow cytometry

THP-1 cells were plated in 12-well culture plates (Corning Costar) as described above. The cells were washed three times with 1X PBS and detached from plates using Accutase (A6964; Sigma-Aldrich) incubation for 15 minutes at 37°C. The detached cells were collected on ice and then centrifuged (settings). Human TruSatin FcX™, FcR Blocking Reagent (1 µg IgG/10^6^ cells in 100 µl staining volume; BioLegend, #422301) was applied to decrease the non-specific binding for 10 minutes on ice. Cells were subsequently stained with Brilliant Violet 785™ anti-human CD14 Antibody (1:1000; BioLegend, #301840), APC anti-human CD86 Antibody (1:1000; BioLegend, #305412), and PE anti-human CD11b Antibody (1:1000; BioLegend, #301306) for 30 minutes in the dark. The cells were then fixed and permeabilized with 1% paraformaldehyde (PFA). Flow cytometry data were acquired on a BD LSRII flow cytometer (BD Biosciences) with FACS Diva software (BD) and analyzed using FlowJo software (FlowJo, LLC).

### Cell lysis and sample preparation for mass spectrometry

The cells for proteomic analysis were cultured, as described above. After the indicated time points of PMA incubation followed by resting, the media was discarded, and the cells were washed thrice with ice cold PBS. The cells were lysed, and harvested using SDS lysis buffer (2% SDS in 50 mM TEABC) tubes and sonicated using a probe sonicator (Branson Digital Sonifier) on ice for 5-10 minutes (20% amplitude, 10 cycles). The lysates were heated at 95°C for 10 minutes, allowed to cool to room temperature, and centrifuged at 12,000 rpm for 10 minutes. The protein concentration in the lysates were estimated by the Bicinchoninic acid (BCA) assay (Pierce, Waltham, MA). 200 µg proteins from each condition were reduced and alkylated with dithiothreitol (DTT) at 60°C for 20 minutes and 20 mM iodoacetamide (IAA) at room temperature for 10 minutes in the dark, respectively. The protein samples were then subjected to acetone precipitation with five volumes of chilled acetone at −20°C for 6 hours. Protein pellets were obtained by centrifugation at 12,000 rpm for 15 minutes at 4 °C and subjected to trypsin digestion with sequencing grade trypsin (1:20) (Sigma Aldrich) overnight at 37 °C.

### Dimethyl-labeling of tryptic peptides

The tryptic peptides obtained from Condition A, B and C were subjected to reductive dimethylation with 4% (vol/vol) formaldehyde (CH_2_O) (Light), (CD_2_O) (Medium) or (^13^CD_2_O) (Heavy) labels respectively. Following this, 4 µl of 0.6 M sodium cyanoborohydride (NaBH_3_CN) was added to the samples to be labeled with light and intermediate labels and 4 µl of 0.6 M sodium cyanoborodeuteride (NaBD_3_CN) to the sample to be heavy labeled respectively. The mixture was incubated for 1 hour at room temperature. The reaction was quenched with 16 µl of 1% ammonia. Finally, 8 µl formic acid was added, and the three differentially labeled samples were pooled and desalted using C_18_ StageTip, evaporated to dryness under vacuum, and subjected to Stage-tip based Strong-cation exchange (SCX) fractionation as described previously (24).

### Mass spectrometry analysis

Mass spectrometric analyses of the SCX fractions were carried out using a Q Exactive HF Hybrid Quadrupole-Orbitrap mass spectrometer (Thermo Fisher Scientific, Bremen, Germany) coupled to Easy-nLC1200 nano-flow UHPLC (Thermo Scientific, Odense, Denmark). The data were acquired for each of the samples in biological quadruplicates. Briefly, tryptic peptides obtained from StageTip-based SCX fractionation were reconstituted in 0.1% formic acid and loaded on a nanoViper 2 cm (3 µm C18 Aq) trap column (Thermo Fisher Scientific). Peptide separation was carried out using EASY-Spray C18 analytical column (50 cm, 75µm PepMap C18, 2 µm C18 Aq) (ES801, Thermo Fisher Scientific) set at 40°C. Peptide separation was carried out at a flow rate of 250 nl/min using a binary solvent system containing solvent A: 0.1% formic acid and solvent B: 0.1% formic acid in 80% acetonitrile. A linear gradient of 5-30% solvent B over 150 minutes, followed by a linear gradient of 30-95% solvent B for 5 minutes, was employed to resolve the peptides. The column was re-equilibrated to 5% solvent B for an additional 20 minutes. The total run time was 180 minutes. Data were acquired in positive mode using a data-dependent acquisition method wherein MS1 survey scans were carried out in 350-1650 m/z range in Orbitrap mass analyzer at a mass resolution of 120,000 mass resolution at 200 m/z. Peptide charge state was set to 2-6, and dynamic exclusion was set to 30 s along with an exclusion width of ± 20 ppm. MS/MS fragmentation was carried out for the most intense precursor ions selected at top speed data-dependent mode with the maximum cycle time of 3 seconds HCD fragmentation mode was employed with a collision energy of 30% and detected at a mass resolution 15,000 at m/z 200. Internal calibration was carried out using a lock mass option (m/z 445.1200025) from ambient air.

### Data analysis

Protein identification and quantification were performed using Proteome Discoverer Version 2.3 with the following parameters: carbamidomethyl of cysteine as a fixed modification, and oxidation of methionine, deamidation of asparagine and glutamine, acetylation (protein N terminus), quantitation labels Dimethyl, Dimethyl:2H4 and Dimethyl:2H(6)13C(2) on N-terminal and/or lysine were set as variable modifications. Trypsin as specified as proteolytic enzyme with maximum of 2 missed cleavages allowed. The searches were conducted using the SequestHT node against the Uniprot-Trembl Human database (v2017-10-25), including common contaminants. Mass accuracy was set to 10 ppm for precursor ions and 0.02 Da for MS/MS data. Identifications were filtered at a 1% false-discovery rate (FDR), accepting a minimum peptide length of 7 amino acids. Quantification of identified proteins referred to the razor and unique peptides and required a minimum ratio count of 2. Dimethyl-based relative ratios were extracted for each protein/condition using the Minora Feature Detector node and were used for downstream analyses.

### Bioinformatics analysis

Protein abundances across multiple replicates were scaled, log-transformed, normalized using the cyclic loess method, and analyzed for differential expression in limma v3.38.3 (25) in R/Bioconductor (v3.5.2, https://www.r-project.org/; v3.8 https://bioconductor.org). The treatment conditions were used for contrasting. Proteins expressed with a log2-fold change ≥±2 were considered as differentially expressed. Volcano plots were drawn using the EnhancedVolcano R package (v 1.0.1) and proteins with –log10 (p-value) ≥ 1.25 were considered to be significant. Heatmaps of expression data and *k*-means clustering was carried out in Morpheus using Euclidean complete linkage (https://software.broadinstitute.org/morpheus/). Significant clusters of genes that were overexpressed in conditions A, B, and C were extracted, and enriched biological processes were identified using Enrichr (https://amp.pharm.mssm.edu/Enrichr/). Hypergeometric enrichment-based pathway analysis was carried out using ReactomePA (1.28.0) in R (v3.6.0, https://www.r-project.org/)/Bioconductor (v3.9 https://bioconductor.org) with clusterProfiler 3.12.0 (26). Genesets with a minimum of 15 genes were considered for the analysis. The plot was visualized using the ggplot2 package (v3.2.1) (https://cran.r-project.org/web/packages/ggplot2). Significantly changing clusters for each condition from the *k*-means clustering analysis were subjected to network analysis using STRING in Cytoscape (version 3.7.1). The network properties were calculated using NetworkAnalyzer and visualized in Cytoscape using betweenness centrality and degree parameters. The entire networks were further subjected to clustering to identify significant sub-clusters using the MCODE app (v1.5.1) (27) in Cytoscape. The parameters used for clustering included degree cutoff of 2, node score cutoff of 0.2, *k*-core of 2, and Max. Depth of 100. Kinome trees were drawn using KinMap (28) (http://kinhub.org/kinmap). Gene lists for functions such as phagocytosis, reactive oxygen species, and inflammasome complex were obtained from the Molecular Signatures Database (MSigDB, v7.0, https://www.gsea-msigdb.org/gsea/msigdb) (29). Protein kinase and phosphatase lists were obtained from previous studies (30-33)

### Western blot analysis

Protein samples were run on pre-cast NuPAGE™ Bis-Tris gels (Invitrogen) with 1 x MOPS buffer (Invitrogen) and transferred on nitrocellulose membranes, using the iBlot®2 Gel Transfer Device (Invitrogen). Membranes were washed in Tris Buffered Saline with 0.1% Tween-X100 (TBS-T) and blocked with TBS-T containing 5% bovine serum albumin (BSA, Sigma-Aldrich). Membranes were incubated with primary antibodies at 4°C overnight. The following primary antibodies were used: GAPDH (1:5000; ab8245; Abcam), anti-IRF3 (1:1000; D83B9; cat#4302S; Abcam), anti-TBK1 (1:1000; cat#3504; Cell Signaling Technology), anti- IL1B (1:1000; cat#12242; Cell Signaling Technology) and SQSTM1 (1:1000; cat#PM045; MBL). Membranes were washed in TBS-T and incubated with secondary antibodies (HRP-conjugated, DAKO) for 1 hour at room temperature in TBS-T containing 1% milk or BSA, developed with SuperSignal West Femto Substrate (Thermo Scientific) and captured with LI-COR Odyssey system (LI-COR Biosciences, Lincoln, NE, USA).

### RNA isolation and quantitative real-time PCR (qPCR) analysis

Undifferentiated THP-1 and PMA-differentiated cells (2 × 10^5^ cells) in biological duplicates per condition were stimulated with TLR agonists- CL075 (TLR8; tlrl-c75; Invivogen, 5 μg/ml), CpG2006 (TLR9; tlrl-2006; Invivogen, 10 μM), Flagellin (TLR5; tlrl-stfla; Invivogen, 100 ng/ml), FSL1 (TLR2/6; tlrl-fsl; Invivogen, 100 ng/ml), LPS 0111:B4 (TLR4; tlrl-3pelps; Invivogen, 200 ng/ml), LPS K12 (TLR4; tlrl-eklps; Invivogen, 200 ng/ml), Pam3CSK4 (TLR1/2; tlrl-pms; Invivogen, 200 ng/ml), Poly (I:C) (TLR3; vac-pic; Invivogen, 10 μg /ml), R837 (TLR7; tlrl-imqs; Invivogen, 10 μg/ml), and R848 (TLR7/8; tlrl-r848; Invivogen, 100 ng/ml) for 4 hours. Post stimulation, total RNA was isolated using RNeasy Mini columns, followed by DNAse digestion (Qiagen), according to the manufacturer’s protocol. The purity and concentrations of RNA were determined using NanoDrop 1000 (Thermo Scientific). cDNA was prepared with High-Capacity RNA-to-cDNA™ (Applied Biosystems). Quantitative real-time PCR (qPCR) analysis was performed on StepOne Plus Real-Time PCR cycler (Thermo Fisher Scientific) using PerfeCTa qPCR FastMix UNG (Quantabio) and FAM Taqman Gene Expression Assays: *IL6* Hs00985639_m1, *IL1B* Hs00174097_m1, *TNF* Hs01113624_g1, *TBP* Hs00427620_m1, *IL8* Hs00174103_m1 (Life Technologies) in 96-well format in technical duplicates. Relative expression compared to the unstimulated control samples and TBP as a house keeping gene was calculated in R 3.3.2 as described previously (25).

### Data availability

Mass spectrometry-derived data have been deposited to the ProteomeXchange Consortium (http://proteomecentral.proteomexchange.org) via the PRIDE partner repository (26) with the dataset identifier: PXD015872.

## Results

### PMA differentiation induces changes in morphology and expression of cell surface markers in THP-1 cells

We aimed to investigate the effects of various concentrations and duration of incubation with PMA on the differentiation of THP-1 monocytic cells to macrophage-like cells. We chose three commonly used differentiation protocols for the analysis based on previous studies (7, 12, 27). Light microscopy analysis revealed changes in PMA induced morphology, including increased cellular adhesion and spread morphology. In concordance with the results observed by Starr *et al.* (28), the changes in cell morphology were dependent on the concentration and the duration of incubation, with cells treated with PMA and rested for two days showing a significant increase in cytoplasmic volume with increased adherence (**Figure 1A**). The differentiation was more pronounced in Condition A in comparison with the conditions B and C. Additionally, flow cytometry analyses indicated an increase in side scatter (SSC), in condition A with respect to conditions B and C. The cells treated with PMA but not rested (Condition C) closely resembled the undifferentiated THP-1 cells in regard to these properties (**Figure 1B**). In addition to the morphological changes, the differential expression of cell surface markers CD86, CD11b (ITGAM), and CD14 were also monitored. CD86, a cell surface glycoprotein expressed on all antigen-presenting cells, was found to be expressed to a similar extent in all the three protocols tested. In concordance with earlier reports, the expressions of CD11b and CD14 were found to be lower in Condition C in comparison to the other two conditions tested. We also observed an effect on the expression based on the duration of incubation and period of rest. The expression of both CD11b and CD14 was comparable in conditions A and B with the highest surface expression observed in condition A (**Figure 1C**). Our data, therefore, confirm the previous findings that the degree of differentiation induced by PMA treatment varies depending on the concentration and period of rest post-PMA treatment.

**Figure 1:**
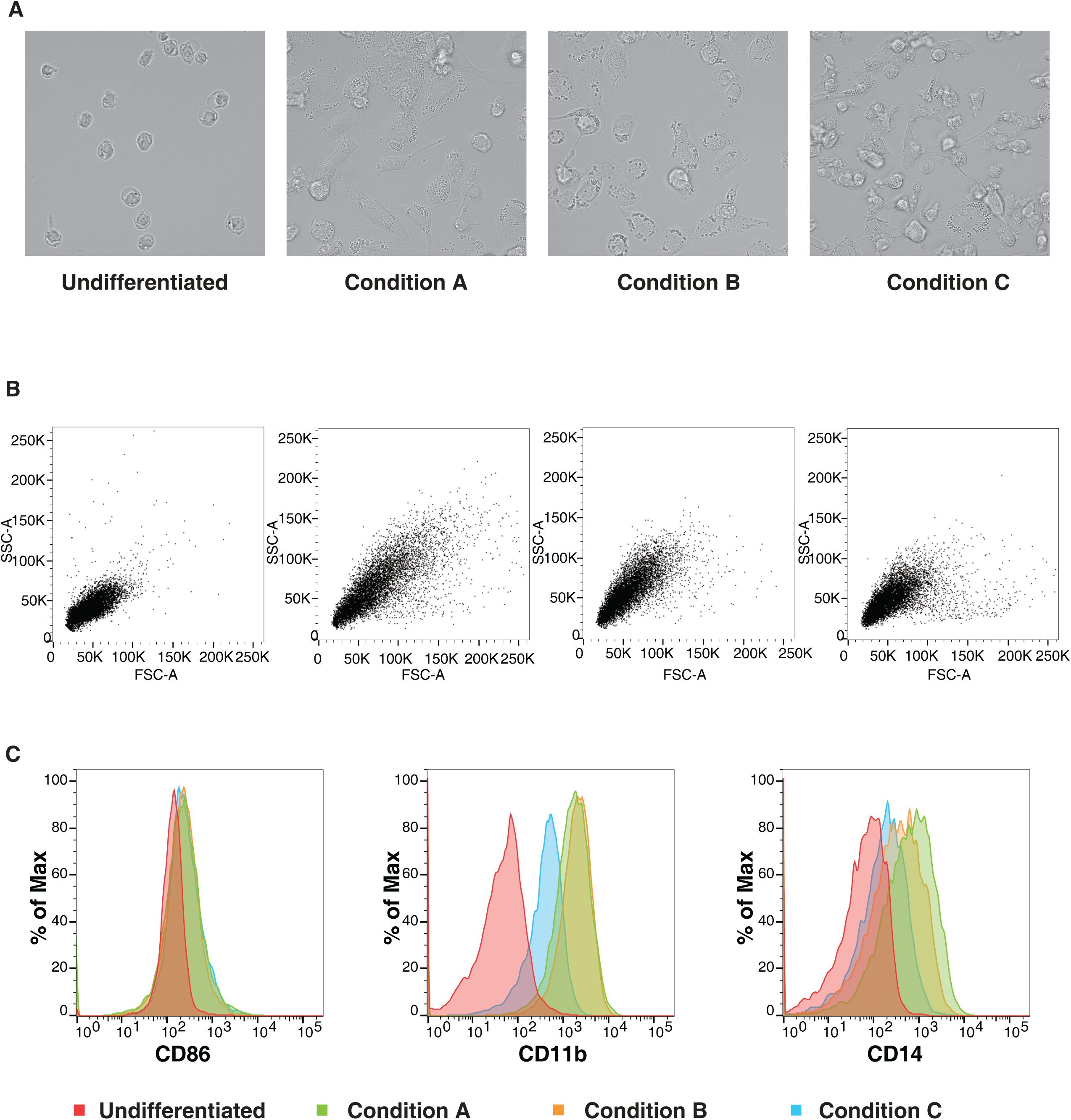
Changes in cell morphology and surface receptor expression are dependent on PMA- mediated differentiation conditions. (A). Representative bright-field images (Scale bar:100 μm) (B) Representative forward and side light scatter plot (C) Representative flow cytometric analysis of THP-1 cells stained using anti- CD86, CD11b or -CD14 of THP-1 cells differentiated with varying concentration of PMA (Condition A: 50ng/ml PMA 72h, +48h rest, Condition B: 50ng/ml PMA overnight, + 48h rest and Condition C: 5 ng/ml PMA 48h, +3h rest). Data is representative of at least three independent experiments

### Quantitative proteomic analysis reveals diverse proteome expression profiles in response to varying differentiation protocols

To evaluate the proteome-wide expression changes upon PMA treatment, we performed quantitative proteomic analysis using a stable isotope dimethyl labeling approach (**Figure 2A**). From four independent biological replicates, 5,277 proteins were identified, of which 5,006 proteins were quantified in at least one replicate. A total of 3,623 proteins were identified and quantified in all four replicates providing a global view of changes in protein expression upon PMA treatment (**Supplementary Table 1**). Principal component analysis (PCA) revealed distinct clustering of each treatment condition with the biological replicates grouped (**Figure 2B**). The highest variance was observed in all replicates of condition C and can likely be explained by the fact that the morphological phenotype observed was much closer to the monocytic cell type rather than the macrophage-like phenotype.

**Figure 2:**
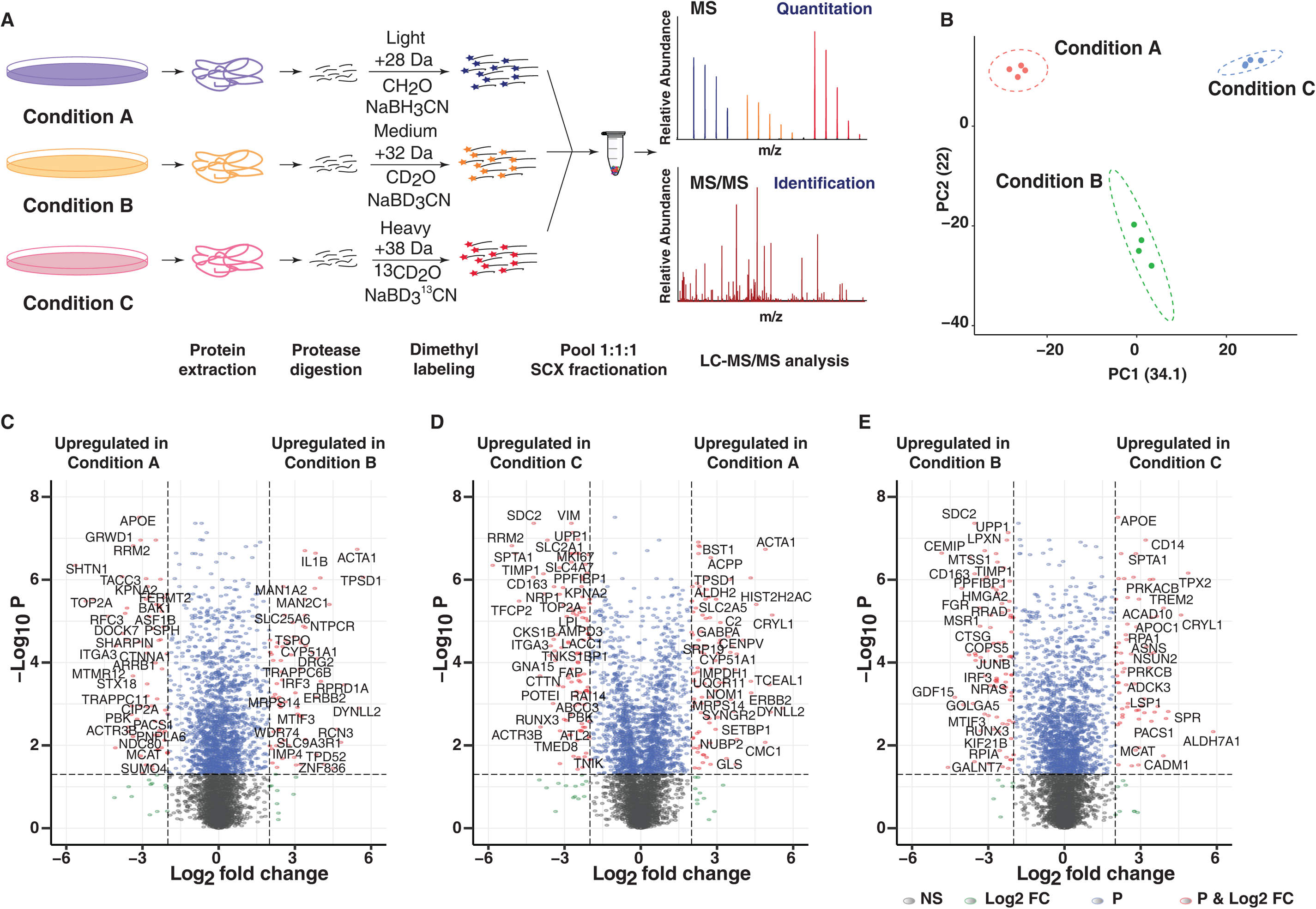
Quantitative proteomic analysis of PMA-induced changes. (A) The experimental strategy employed for comparative proteome analysis in response to varying PMA-mediated differentiation protocols. Dimethyl labeling-based quantitative proteomic approach was employed to identify and quantify the proteome changes in Condition A (Light), Condition B (Medium), and Condition C (Heavy).The data were acquired in biological quadruplicates and only proteins identified and quantified in all 4 replicates were considered for further analysis. (B) Principal component analysis (PCA) reveals that the three treatment conditions for monocyte-to macrophage differentiation of THP-1 cells segregate from each on the basis of concentration and duration of PMA treatment. All replicates of a given condition cluster together suggesting minimal biological variability. (C-E) Volcano plot displaying differential expressed proteins between (A) Condition A vs. B (B) Condition C vs. A and (C) Condition C vs. B. The vertical axis (y-axis) corresponds to the mean expression value of log 10 (q-value), and the horizontal axis (x-axis) displays the log 2-fold change value. The red dots represent overexpressed proteins, and the green dots represent proteins with downregulated expression. Positive x-values represent overexpression, and negative x-values represent down-regulation.

Next, we applied a log2 (fold-change) and adjusted p-value cutoff of 2 and < 0.05 resulting in the identification of 324, 415 and 413 proteins upregulated and 299, 321 and 338 proteins downregulated in condition B with respect to condition A (B/A), condition C with respect to condition A (C/A) and condition B (C/B), respectively (**Figure 2 C-E, Supplementary Figure 1A**). The segregation of condition A from condition B and C was mainly driven by differential expression of several proteins involved in vesicle-mediated transport (KIF13B, KIF2C, KIFC1, STX18), cell cycle regulation and mitosis (RRM1 and RRM2), DNA polymerase complex subunits- POLD1 and POLD2, topoisomerase TOP2A, TACC3, and PRKACB. Interestingly, proteins involved in innate immune response such as TBK1, IRF3, IL1B, scavenger receptor SCARB1, and mitochondrial translation initiation factor MTIF3 were significantly differentially expressed in condition B in comparison to conditions A and C. On the contrary, members of the aldehyde dehydrogenase family such as ALDH1L2, ALDH2, members of serine/threonine-protein kinase C (PRKCA, PRKCB, PRKCD), proteins involved in amino acid metabolism-GOT1, PSAT1, ASNS among others were found to be significantly upregulated in condition C in comparison to conditions A and B. It has been previously shown that PMA-mediated differentiation increases the expression of PKC isoenzymes albeit to a varied extent (29). The results from the MS analysis were further confirmed by validating the expression dynamics of select proteins using immunoblot analysis. Consistent with MS results, the expression of IRF3, TBK1, IL1B, and SQSTM1 were upregulated in cells differentiated for 3 and 5 days (**Supplementary Figure 1B**).

Monocyte-to-macrophage differentiation is reportedly associated with changes in the expression of cell surface proteins, and this phenomenon has been utilized to distinguish macrophage subtypes by their pattern of cell surface receptor expression (7, 30). Our analysis revealed increased expression of known macrophage cell-surface markers such as TFRC (CD71), FCGR1B, scavenger receptors- CD163, MSR1, and SCARB2 mainly in conditions A and B. Notably, CD163, a member of the scavenger receptor cysteine-rich (SRCR) superfamily class B, has been previously reported to be highly expressed in macrophages with low expression reported in monocytes, dendritic cells and Langerhans cells (31, 32). On the contrary, the expression of CD68 and CD14 in agreement with previous studies were downregulated in condition B, suggesting a phenotype closer to macrophages. Interestingly, the expression of CD36 and ITGAM (CD11b) was upregulated in condition A with respect to conditions B and C, whereas that of CD68 was downregulated in both conditions A and B with respect to condition C (**Supplementary Figure 1C, Table 1**). Comparison with the known macrophage differentiation markers (GO:0030225) revealed significant induction of CSF1R, TLR2, APP, RB1, CASP8, FADD, MMP9 and ICAM1 in conditions A and B with the exception of transcription factor PU.1 (SPI1) and PRKCA that were significantly up in condition C and CDC42 that was expressed to similar extent in all three protocols tested.

### Cluster analysis reveals differentiation protocol-specific regulation of cellular processes and signaling pathways

To gain further insights into specific expression profiles, we performed unsupervised hierarchical clustering of proteins quantified, which revealed a high correlation between biological replicates. Using Euclidean average and k-means clustering, the differentially expressed genes were segregated into 10 major clusters (**Figure 3A, Supplementary Table 2**). Cluster 1, 7, and 10 included proteins upregulated in conditions B, A, and C, respectively. Cluster 3 includes 290 proteins that were expressed to a similar extent in conditions B and C but downregulated in condition A. On the contrary; Cluster 6 included 458 proteins that were expressed to a similar extent in conditions A and B but downregulated in condition C and cluster 9 comprised of proteins overexpressed in conditions A and C with respect to condition B, but with an overall increased expression observed in all replicates of condition C. Clusters 2, 4 and 5 showed similar expression of proteins across all three protocols tested with varying expression across replicates. Altogether, our analysis demonstrates the existence of common and differentiation-protocol-specific proteome signatures.

**Figure 3:**
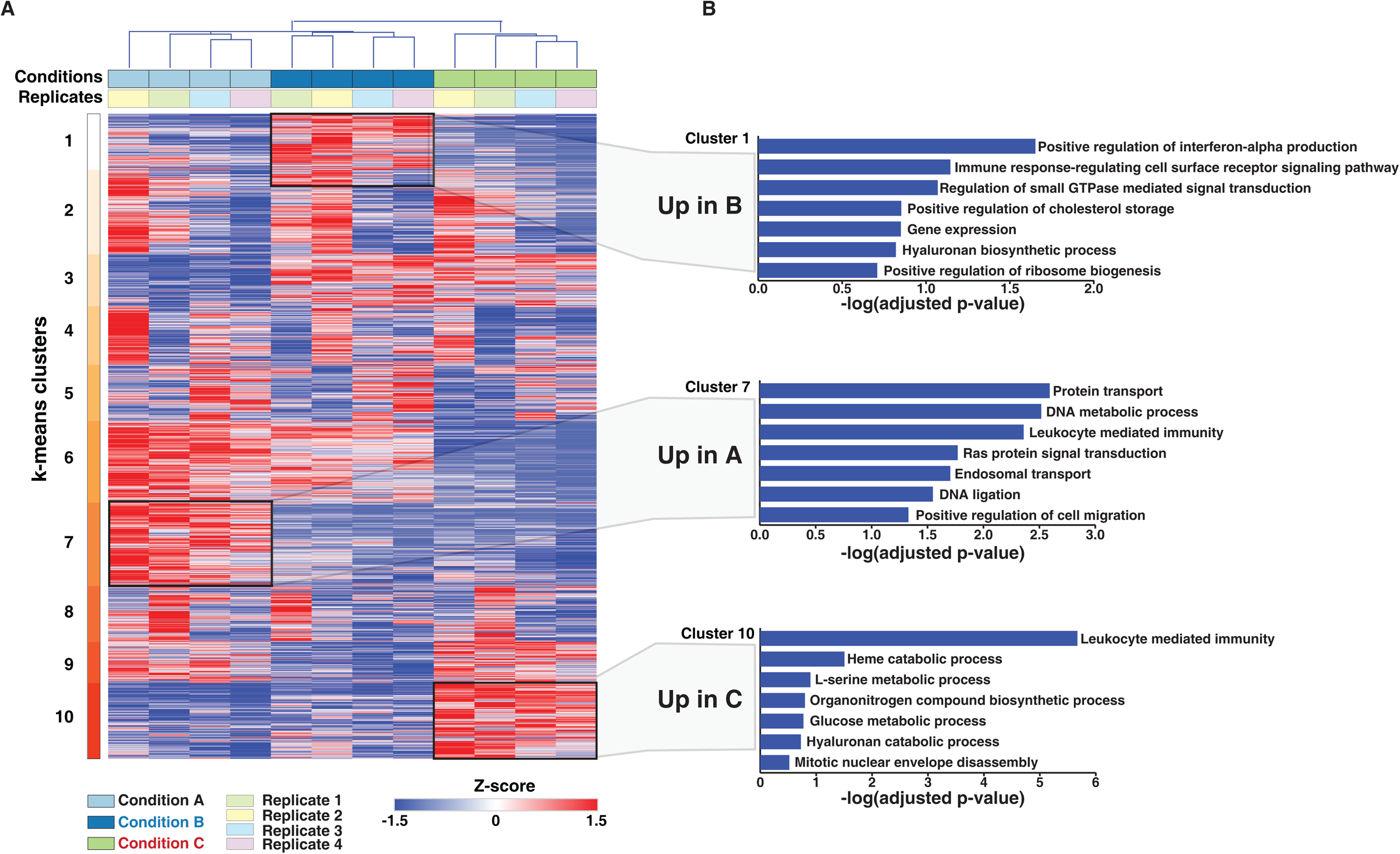
Protein expression dynamics analysis upon PMA induced monocyte to macrophage differentiation. (A) Protein expression patterns in response to differentiation protocols A, B, and C were analyzed. Log2 transformed, z-score normalized and scaled expression of proteins identified and quantified in all replicated were plotted, and k-means clustering was carried out. K-means clusters 1, 7 and 9 which showed overexpressed proteins exclusive to conditions A, B, and C, have been highlighted. (B) The proteins exclusive to each condition were subjected to Gene Ontology (GO) analysis using Enrichr to understand their function. Selected significantly enriched GO terms (biological processes) (p-value⩽0.005) have been highlighted.

Gene ontology analysis of the differentiation protocol-specific clusters (clusters 1, 7, and 10) revealed significant enrichment of several biological processes (adjusted p-value < 0.05) (**Figure 3B, Supplementary Table 3-5**). Across the three differentiation conditions, several metabolic processes were enriched. Processes indicative of differentiation, such as cell migration and mitotic nuclear envelope assembly, were also enriched. While immune response-regulating cell surface receptor signaling pathway was observed to be primarily enriched in condition B, the process of leukocyte mediated immunity was found to be enriched in conditions A and C (**Supplementary Figure 2**). Regulation of translation and gene expression were among the significantly enriched processes in Cluster 3 and 6, whereas in the case of Cluster 9, which included proteins expressed to a similar extent in conditions A and C but downregulated in condition B, we observed enrichment of carbohydrate and fatty acid metabolic processes. (**Supplementary Figure 2**). The findings described above are consistent with the morphological changes observed with processes indicative of differentiation, such as cell migration, gene expression, ribosome biogenesis as well as that of immune cell function which were largely enriched in conditions A and B respectively.

### Pathway enrichment and network analysis reveals kinases as key regulatory hubs of the PMA mediated differentiation processes

We next aimed to delineate the signaling pathways affected during monocyte-to-macrophage differentiation. Pathway enrichment analysis using the Reactome database revealed differentiation protocols specific enrichment of signaling pathways such as VEGF signaling pathway (enriched in conditions A and B), metabolism of nucleotides and porphyrins (enriched in condition A), Fc epsilon R1 signaling (enriched in condition B), activation of NADPH oxidases by RhoGTPases, and glycosphingolipid metabolism (enriched specifically in condition C) (**Figure 4A, Supplementary Table 6**). Interestingly, cell surface signaling mediated by ephrins, integrin, semaphorin, and syndecan interactions were significantly enriched in condition A with respect to condition C with no apparent differences observed with condition B. On the contrary, signaling pathways involved in clearing infections and immune responses were significantly enriched in condition B. Apoptotic pathways were found to be downregulated, whereas transamination and amino acid synthesis were upregulated in condition C.

**Figure 4:**
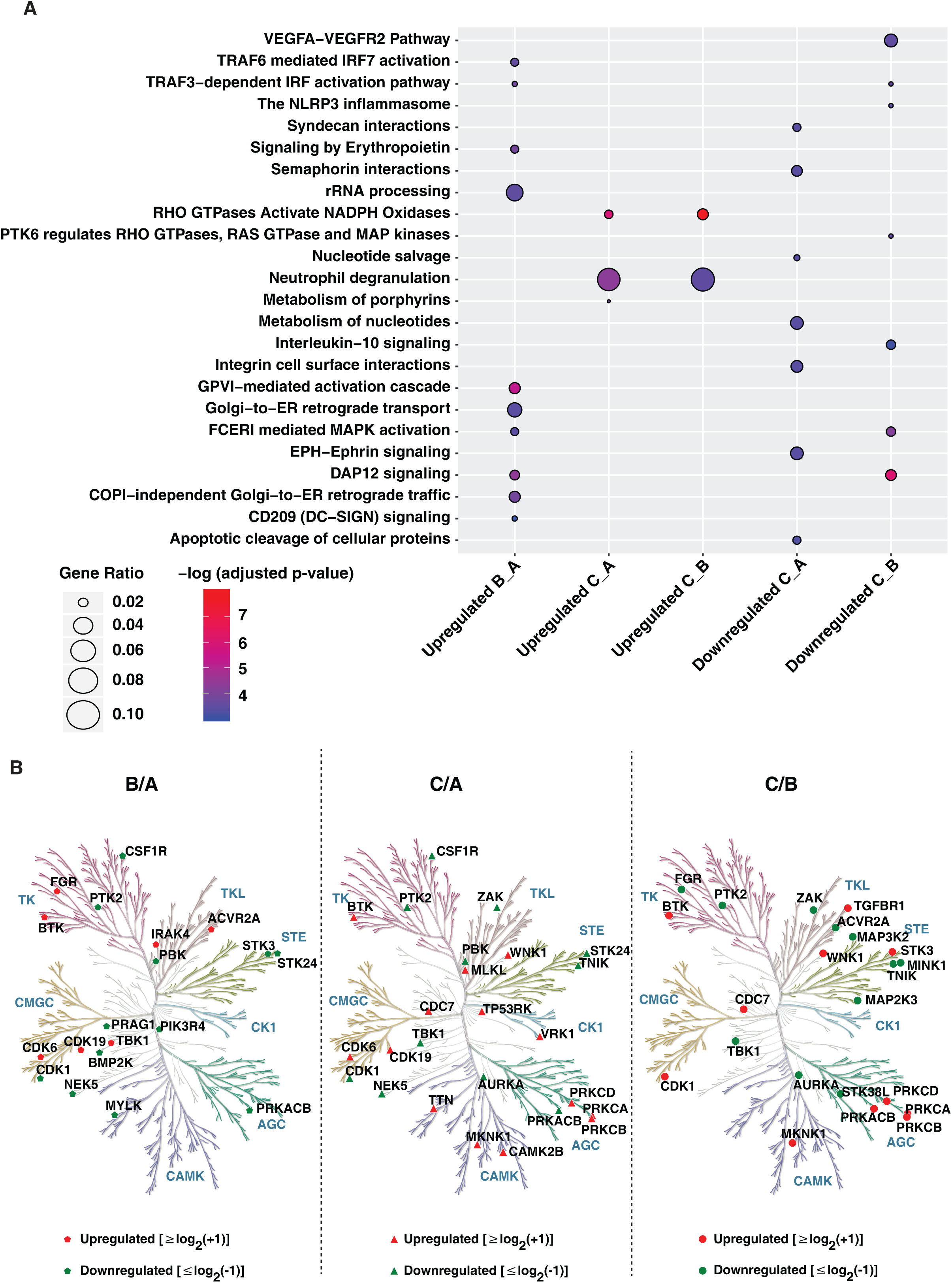
Gene ontology analysis of proteins upon PMA induced monocyte to macrophage differentiation. (A) Gene Ontology analysis of upregulated and downregulated proteins in conditions A, B, and C using ReactomePA. (B). Kinome trees showing differential regulation of protein kinases in response to differentiation conditions A, B, and C. The phylogenetic kinase relationship (Manning et al., 2002) were generated using KinMap. Protein kinases identified and quantified in our study are indicated as gray and black circles, respectively.

Network analyses of k-means clusters upregulated in each condition **(Supplementary Figure 3-5, Supplementary Table 7)** enabled identification of distinct hub proteins. Interestingly, several kinases involved in cellular processes namely cell proliferation, differentiation and regulation of microtubule dynamics such as MAPK1, CDK1, and PRKACB were found to have a high degree of betweenness centrality and exist as key regulatory hubs in the network for condition A. Other key regulatory hubs included DUT, FN1 a glycoprotein involved in cell adhesion and migration processes and SEC13, a core component of the COPII-coated vesicles and nuclear core complex. While proteins involved in innate immune response such as ISG15, IL1B, LYN, COPS5, TBK1, MRPL3, and KRAS were found to be critical regulatory hubs in condition B, enzymes involved in amino acid metabolism such as GOT1, PSAT1, AARS, CBS, IMPDH1; subunit of RNA polymerase II (POLR21), plasma membrane-associated Rho GTPase RAC1 and PRKCD were found to critical regulatory proteins in condition C. Interestingly, a previous study exploring the role of kinases in monocyte-macrophage differentiation observed a pronounced decrease in the expression of regulatory kinases such as CDK1 involved in cell cycle entry and checkpoint in PMA-differentiated THP-1 macrophage-like cells (22). It is well known that PMA activates protein kinase C (PKC) and promotes leukocyte adhesion and migration, and therefore identifying PRKCD as one of the regulatory hubs is indicative of the monocyte-to-macrophage differentiation process.

Further investigation on the effect of PMA differentiation protocols on the extent of expression of other protein kinases as well as phosphatases revealed increased expression of several kinases belonging to diverse classes (**Figure 4B**). Notably, protein tyrosine kinases such as CSF1R and PTK2 was overexpressed in condition A, FGR, a member of the c-Src family tyrosine kinases known to be induced by PMA (33-35) in condition B, and BTK2 in condition C, respectively. Interestingly, increased expression of several kinases was observed in condition C including members of the protein kinase C family-PRKCB, PRKCD which are known markers of immune cell differentiation and inflammation, MAP kinase-interacting serine/threonine-protein kinase 1 (MKNK1), MAPK14 and WNK1, a known regulator of ion transport proteins involved in the differentiation and migration of endometrial stromal cells (36) and glioma cells (37). Among the protein phosphatases, 80 were identified and quantified in our dataset, with a vast majority expressed to a similar extent in all conditions tested. Of note, phosphatases belonging to the HP2 family namely: prostatic acid phosphatase ACPP and acid phosphatase 2 (ACP2), were significantly dysregulated in expression with ACPP over 20-fold overexpressed in condition C with similar level of expression observed in conditions A and B. On the contrary, ACP2, a lysosomal acid phosphatase was overexpressed 2-fold higher in condition A compared to both condition B and C. Among the dual specificity protein phosphatases (DUSPs), SSH3, a member of the slingshot family was upregulated in condition C. SSH3 is known to specifically dephosphorylate and activate Cofilin, one of the key regulators of actin filament dynamics and remodeling (38). Of the 16 protein tyrosine phosphatases (PTPs) identified, the expression of PTPN7, PTPRC were high in condition C whereas PTPN12, PTPN23, PTPN2, PTPRA, PTPRK, and PTPRU were overexpressed in condition A. Interestingly, PTPRE and to a smaller extent, PTPN6 demonstrated increased expression in condition B. Collectively, the protein kinases and phosphatases identified in this study indicate their differential capacity in the regulation of microtubule stability, actin cytoskeleton reorganization and autophagy in macrophages which in turn are required for their functional responses including phagocytosis, antigen presentation, DAMP and PAMP mediated immune signaling.

### PMA-induced monocyte-to macrophage differentiation modulates the expression dynamics proteins involved in innate immune signaling

Several previous studies have used THP-1 cells as a model for the immune modulation approach; therefore, we assessed the impact of differentiation protocols on the expression of proteins involved in innate immune signaling. Analysis of the expression profile of proteins involved in Toll-like receptor (TLR) signaling pathways identified several known downstream effector proteins (**Figure 5A, Supplementary Table 8**). A majority were expressed at similar levels in all three conditions, suggesting that the effect of following secondary inflammatory stimuli would be mostly independent of PMA stimulation. Among the TLR receptors, we identified only TLR2 with over 2-fold expression in conditions A and B. TLR4 co-receptors, namely CD14 and CD180, were found to be expressed relatively lower extent in conditions A and B. CD14 also known as monocyte marker is downregulated upon differentiation. The other signaling regulators and adaptor proteins such as CNPY3 and UN93B1 and CD36 were in general upregulated in condition A in comparison to conditions B and C.

**Figure 5:**
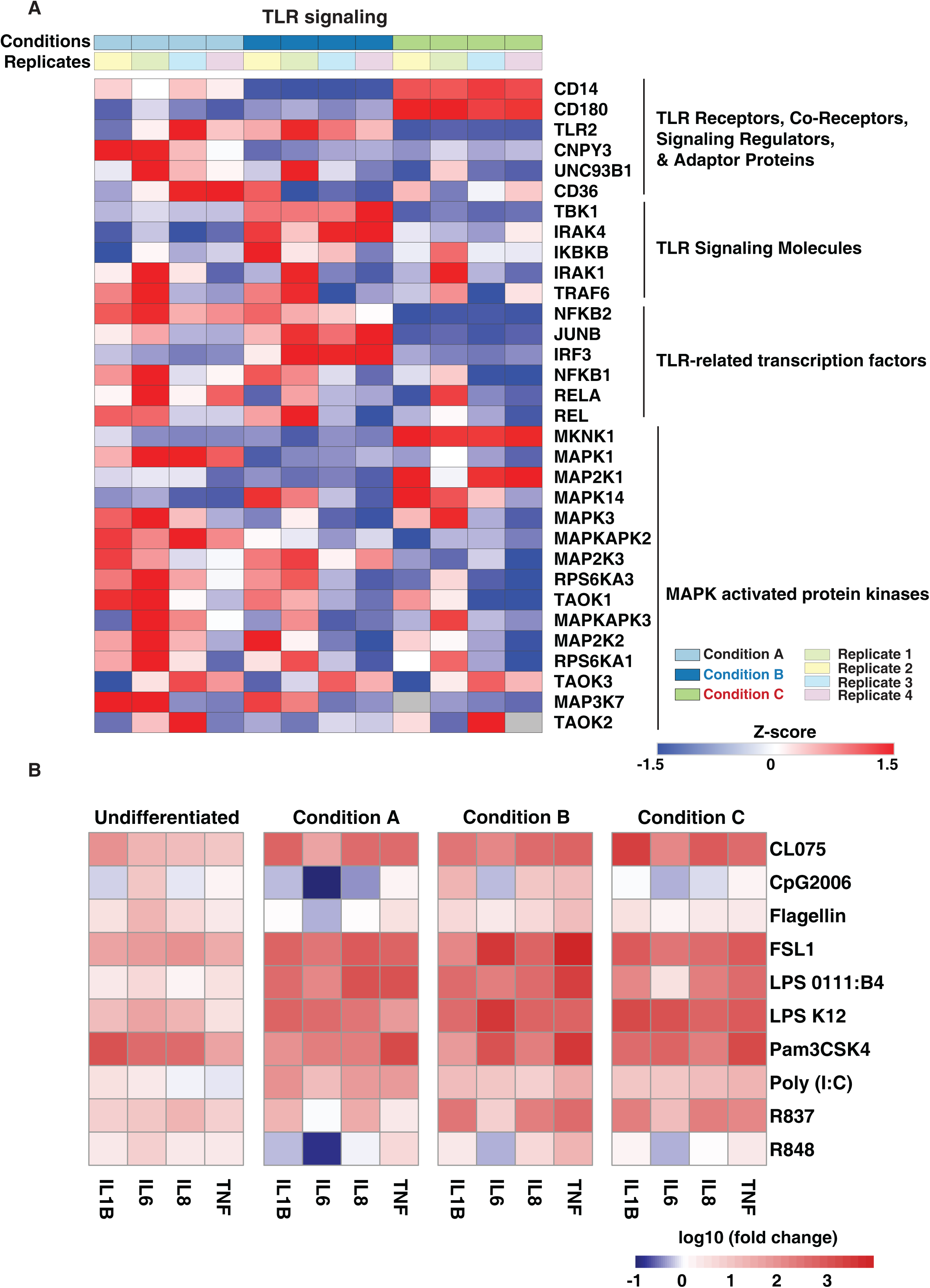
Expression dynamics of genes involved in innate immune signaling in response to PMA. (A) Heatmap showing the proteome fold change of the subset of genes involved in innate immune signaling. Scale indicates the level of expression (Log2-expression values, z-transformed, scaled). (B) Dynamic expression pattern to assess the expression levels of pro-inflammatory cytokine mRNA induced by TLR agonists: CL075-TLR7/8, CpG2006-TLR9, Flagellin-TLR5, FSL1- TLR2/6, LPS 0111:B4, and LPS K12 – TLR4, Pam3CSK4-TLR2/1, Poly I:C-TLR3, R837-TLR7, and R848-TLR7/8. Heatmap depicts Log10 fold change of normalized gene expression for pairwise comparisons of mRNA levels.

Signaling mediated by TLRs requires the assembly of the Myddosome complex. Although we identified Myd88 (39), quantitation was obtained in only conditions A and B. The other components including IRAK1, IRAK4 and TRAF6 were overexpressed in condition B with slightly increased expression in condition A compared to condition C. Of the transcription factors essential for mediating TLR activity, we observed increased expression in mostly conditions A and B with JUNB, highly expressed in condition B. RelB, a member of NF-κB family, was identified in only conditions A and B (40, 41). The expression of Interferon regulatory transcription factors also varied across the conditions tested with over 4-fold expression IRF3 expression in condition B in comparison to conditions A and C whereas IRF5, a key regulator of antiviral immune response, was 2-fold overexpressed in condition C with respect to conditions A and B.This indicates that using condition C to study TLR signaling and/or non-canonical NF-κB signaling pathway may likely alter the outcome of the experiment, and the subsequent phenotype will mostly be dependent on the PMA stimulation. The expression of kinases belonging to the MAPK family, as described in the earlier section, was generally higher in conditions A and B with the exception of MKNK1, MAP2K1, and MAPK14 which were observed to be upregulated in condition C (**Figure 5A**). Our results are in concordance with a previous report suggesting rewiring of MAPK signaling cascade upon THP1 differentiation (22). We also identified 5 proteins known to be a part of the inflammasome complex that were expressed at similar levels except for NLRP3 that was found to be expressed only in condition B (**Supplementary Figure 6**). We also observed differential expression of proteins known to be involved in the process of phagocytosis and oxidative stress such as FGR, LYN, and pro-inflammatory cytokine IL1B among others which were selectively upregulated in condition B. However, neutrophil cytosolic factors NCF2 and NCF4-regulatory components of the superoxide-producing phagocyte NADPH-oxidase, PYCARD, ITGAL, protein kinase C delta (PRKCD), PTPRC, were found to be upregulated in condition C (**Supplementary Figure 6**). Taken together, our analysis provides insights on the differential expression of immune signaling mediators, transcription factors and effectors that can determine the outcome of the signaling responses following the type of differentiation protocol employed.

The response of activated macrophages by various stimuli involves the secretion of cytokines such as IL-1β, IL-6, IL-8, and TNF-α as a significant component of the innate immune response (42). We, therefore, studied the functional properties, cytokine gene expression in undifferentiated (monocytic), and differentially differentiated THP1 cells (macrophage-like) and compared the dissimilarity and similarity of expression trends among conditions. To assess the changes in the cytokine mRNA expression levels, cells were treated with various TLR ligands: CL075-TLR7/8, CpG2006-TLR9, Flagellin-TLR5, FSL1-TLR2/6, LPS 0111:B4, and LPS K12 – TLR4, Pam3CSK4-TLR2/1, Poly I:C-TLR3, R837-TLR7, and R848-TLR7/8 (**Figure 5B**). Monocytes are the first cells that encounter pathogens and promptly adapt to the environmental situations by regulating the expression of genes activated by the inflammasome, such as IL-1β (43). As anticipated, the TLR2 ligand Pam3CSK4 stimulated undifferentiated cells showed significant cytokine induction of *IL1β* mRNA. Interestingly, it also induced a high level of *IL6* and *IL8*. On the other hand, the average cytokine expression appeared to increase in PMA differentiated cells with respect to the undifferentiated cells, except when it was simulated with CpG2006, Flagellin, and R848. Expression of cytokines upon stimulation with FSL1, LPS 0111: B4, LPS K12, and Pam3CSK4 did not significantly vary among the different differentiation conditions except for *IL6* mRNA which was markedly downregulated upon CpG2006, Flagellin and R848 stimulation in condition A. Overall, the most robust upregulation, especially with respect to *IL6* and *TNF-α* expression levels were observed in condition B (**Figure 5B**). Together these experiments indicate that depending on the context of the experimental question, careful consideration of the differentiation protocols selection must be made to avoid undesired outcomes.

## Discussion

Human monocytic THP-1 cells are extensively used as a model system to study monocyte/macrophage functions. However, to be used as an *in vitro* model mimicking human macrophages, THP-1 cells have to be differentiated, and several protocols have been tested (7, 9, 29, 44). Among these, PMA is most often employed to induce differentiation, with almost similar phenotypes reported in terms of cell morphology, expression of macrophage surface markers, and cytokine production (28, 45). However, the amount and duration of incubation with PMA vary widely across the literature. Additionally, studies demonstrate altered sensitivity, as well as undesirable gene regulation in PMA-differentiated macrophages that may contribute to their differential response to secondary stimuli (12, 28). Although this may likely be the case, one inherent limitation of most of these studies is that their focus has primarily been on characterizing either the transcriptional profile or surface receptor expression (11, 30). Since the effect of any stimuli is a global response, it is imperative to study the impact of differentiation at the global proteome level.

To illustrate changes in the proteome expression dynamics during differentiation of THP-1 cells into macrophage-like cells, we performed a global quantitative proteomic analysis by comparing three established and widely employed PMA-mediated differentiation protocols. Our analysis indicates that in the course of monocyte to macrophage differentiation, the proteome profiles across the three tested protocols display common and differentiation-protocol-specific regulation with the expression of known macrophage differentiation markers significantly induced in two of the three conditions. We also delineated the biological and cellular processes that are impacted to a considerable extent depending on the type of treatment conditions. Importantly, regulation of immune signaling response, protein transport, and regulation of cell migration, all indicative of macrophage function were significantly enriched in conditions A and B. These treatment conditions include the use of a higher concentration of PMA, followed by a period of rest appeared to correlate well with macrophage-like phenotype, unlike what was suggested previously (12).

GM-CSF-mediated differentiation of primary human monocytes to macrophages upregulates the macrophage-specific surface markers/receptors, antigen-presenting function, phagocytosis, anti-microbial activity, lipid metabolism and production of growth factors and pro-inflammatory cytokines as evidenced by the global transcriptome analysis of GM-CSF-induced macrophages (46). We too observed a similar trend with treatment condition B showing a better representation of proteins involved in innate immune signaling while cathepsins-proteases involved in innate immune responses and protein kinases involved in the processes of cell proliferation and differentiation (47, 48) were found to be significantly induced in condition A. The increased expression of *IL1B* upon PMA differentiation has been reported earlier at the transcriptome level (11). Although we did not examine the phagocytic activity of differentiated macrophages in this study, we provide evidence of differential expression of proteins involved in the process of phagocytosis across the three tested protocols. Importantly, proteins including kinases, members of Rho GTPase family and adaptors involved in the formation of lamellopodia and actin cytoskeleton reorganization among others were found to be upregulated in conditions A and B (49). On the contrary, regulatory components of the superoxide-producing phagocyte NADPH-oxidase were upregulated in condition C. Our analysis also resulted in the identification of several integrins, known to be involved in the processes of cell adhesion and cell surface receptor-mediated signaling required to maintain homeostasis as well as host defense and inflammation (50). Although the extent of expression of integrins were almost similar, differential induction of CD49c (ITGA3) and ITGB1 (CD29) (50), were observed in condition A with respect to others. ITGAL (CD11a), a pan-leukocyte marker was found to be upregulated in condition C compared to the other two tested protocols indicating that the differential expression may likely contribute to differences in the initiation of primary immune response (50).

Our analysis further highlights differentiation protocol-specific subsets with several protein kinases serving as central hubs in mediating the differentiation process. This is of vital importance as cellular signaling is largely governed by the regulation of expression of kinases and phosphatases in a cell type-specific manner or upon cell activation (51-53). One of the major findings of this study is the differential expression of key regulatory kinases implicated in cell cycle regulation including cyclin-dependent kinases, NEK family of serine-threonine kinases as well as cAMP/PKA-induced signaling that are collectively responsible for enrichment of signaling pathways involved in differentiation, maturation, and regulation of actin cytoskeleton dynamics (54, 55). Interestingly, a pronounced decrease in the expression of vital regulatory kinases in cell cycle entry and checkpoint including the cyclin-dependent kinases has been reported in a previous study exploring the role of kinases in monocyte-macrophage differentiation (22). Similarly, the members of protein kinase C family of serine- and threonine kinases are known to phosphorylate a wide range of proteins involved in cell maintenance and also serve as receptors for phorbol esters (56). We observed several members of this family to be significantly overexpressed in condition C with respect to the other conditions. The observed effect may be attributed to these cells being in a transition state and therefore, do not demarcate as monocyte/macrophage populations. Results from previous studies also indicate that the monocyte to macrophage differentiation is accompanied by extensive rewiring of the MAPK-signaling cascades (22). This was partly observed in our analysis as well with MAPK1 identified as a key regulatory hub in condition A. As reported earlier, we too observed a moderate increase in the expression of MAP2K3, MAP3K2 and MAP3K7 in conditions A and B in comparison to condition C, thereby confirming that these two protocols induced the differentiation to macrophage-like cells (22).

Cytokine responses induced in monocytes and macrophages are known to vary depending on the stimulus and the type of differentiated macrophage studied. Overall, we observed an increase transcriptional response of pro-inflammatory cytokines upon stimulation with TLR ligands except for R848- a TLR8 agonist and CPG2006, a TLR9 agonist, which showed an overall decrease in the expression of cytokine mRNA expression especially *IL6* mRNA in condition A and C respectively. Between the three different protocols, no apparent differences in the induction of cytokines were observed in the case of flagellin (TLR5 agonist), poly I:C (TLR3 agonist), and LPS (TLR4 agonist). The consensus on the induction of *TNFα* by LPS in monocytes and macrophages remains unclear as some studies report higher induction in monocytes, whereas others report the same in macrophages, and these differences may likely be attributed to the differentiation protocols and purity and type of the ligands. Using rough and smooth LPS, we demonstrate an increased induction of all pro-inflammatory cytokines in all the three protocols tested with sustained increased expression observed in condition B. We also noted that the more differentiated macrophage-like cells had higher *TNFα* inducible responses to the TLR2 agonist Pam3CSK4. On the contrary, the induction of *IL1β* by Pam3CSK4 was downregulated with respect to undifferentiated monocytes in all the three protocols tested. Taken together, our findings suggest that TLR ligands induce comparable levels of cytokine gene expression across the protocols tested with condition B.

In conclusion, conditions A and B were found to express the most number of macrophage markers, and desirable characteristics such as increased expression of cell surface receptors and differentiation markers, increased expression of proteins involved in innate immune signaling, and overall similar responses to TLR ligands. This suggests that the extended duration of rest post PMA treatment may not directly contribute to significant alterations in the proteome expression. Although both expressed a distinct set of proteins, a vast majority of cellular processes, including metabolic responses, were mostly unaffected, suggesting that metabolic reprogramming effects observed during secondary stimuli would be independent of the PMA response. This suggests that of the tested conditions, condition B is the most optimal differentiation protocol to studying innate immune signaling. Minor deviations in the protocols can have unintended effects on the overall experimental setup and subsequently on the results. Therefore, validation of the model solely by means of morphological or cell surface expression of select markers may not suffice and should be supplemented with orthogonal experiments that provide a more significant overview of the cellular state. The present datasets, in particular, the quantitative differences in the proteome repertoire of molecules involved in innate immune signaling, represent a valuable resource to understand and modulate the functionality of monocyte-derived macrophages.

## Acknowledgments

LC-MS data acquisition was performed at the Proteomics and Modomics Experimental Core Facility (PROMEC), Norwegian University of Science and Technology (NTNU). PROMEC is funded by the Faculty of Medicine and Health Sciences at NTNU and the Central Norway Regional Health Authority. Data storage and handling are supported under the NIRD/Notur project NN9036K. We thank Lars Hagen, Animesh Sharma, and Aditya Kumar Sharma for their technical assistance and Geir Slupphaug for his continued support.

## Funding

This research was funded by the Research Council of Norway (FRIMEDBIO “Young Research Talent” Grant 263168 to R.K.K.; and Centres of Excellence Funding Scheme Project 223255/F50 to CEMIR), Onsager fellowship from NTNU (to R.K.K.), and Regional Health Authority of Central Norway (90414000 to KB).

## Abbreviations

BCA: Bicinchoninic Acid
DTT: Dithiothreitol
FACS: Fluorescence-Activated Cell Sorting
GO: Gene Ontology
HCD: Higher-energy Collisional Dissociation
IAA: Iodoacetamide
MS/MS: Tandem mass spectrometry
PCA: Principal Component Analysis
PMA: Phorbol-12-myristate-13-acetate
SCX: Strong Cation Exchange
TLR: Toll-Like Receptor
UHPLC: Ultra-High-Performance Liquid Chromatography

