## Supplementary Figure 1 for "Dose-dependent phorbol 12-myristate-13-acetate-mediated monocyte-to-macrophage differentiation induces unique proteomic signatures in THP-1 cells"

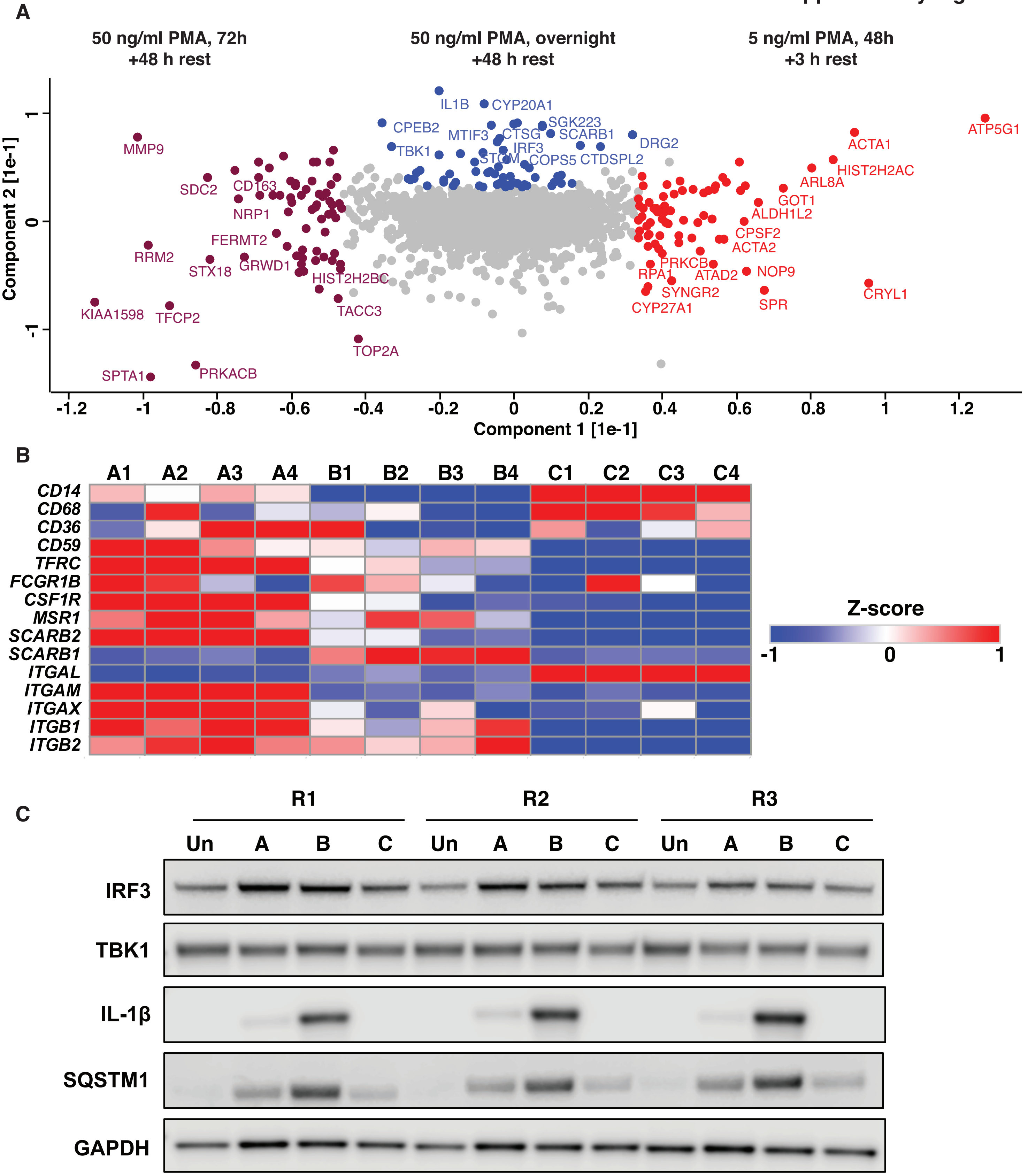


**Supplementary Figure 1**

(A) Principal component analysis (PCA) based on the proteins driving the segregation between the monocyte-to-macrophage differentiation protocols. Proteins highlighted in blue, purple, and red segregate conditions B, A and C, respectively.

(B) Heatmap depicting cell surface receptor expression across the three differentiation protocols. Scale indicates the level of expression (Log2-expression values, z-transformed, scaled.

(C) Western blot analysis of THP-1 cells differentiated with various concentrations of PMA in comparison with undifferentiated THP-1 monocytes using antibodies against IRF3, TBK1, IL1B, and SQSTM1. Immunoblot analysis was performed in triplicates. GAPDH was used as a loading control.
