## Supplementary Figure 2 for "Dose-dependent phorbol 12-myristate-13-acetate-mediated monocyte-to-macrophage differentiation induces unique proteomic signatures in THP-1 cells"

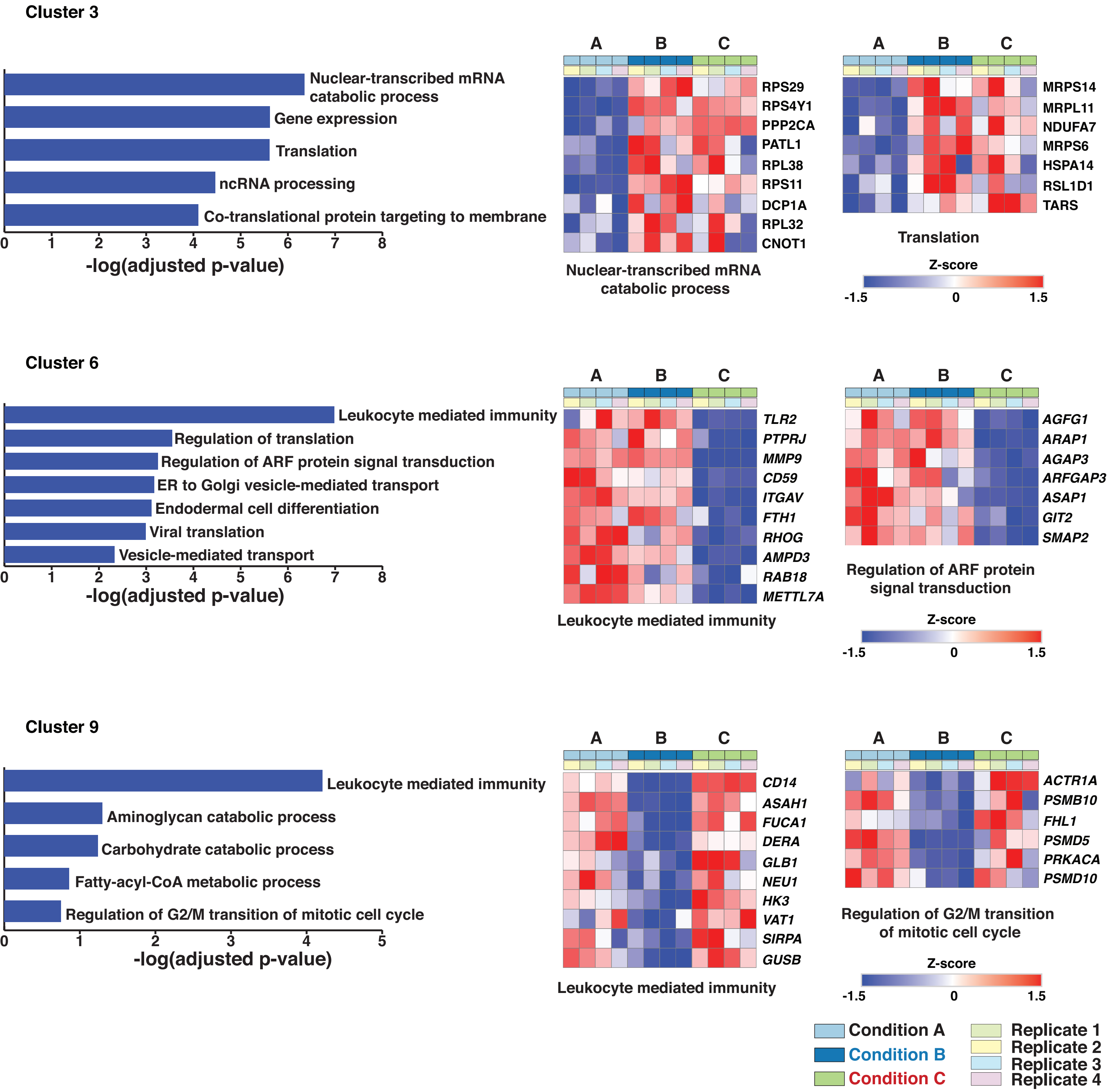


**Supplementary Figure 2**

Significantly enriched biological processes (p-value ≤ 0.005) for K-means clusters 3, 6 and 9. Heatmap depicts the expression changes of selected genes associated with the respective biological process.
