## Supplementary Figure 3 for "Dose-dependent phorbol 12-myristate-13-acetate-mediated monocyte-to-macrophage differentiation induces unique proteomic signatures in THP-1 cells"

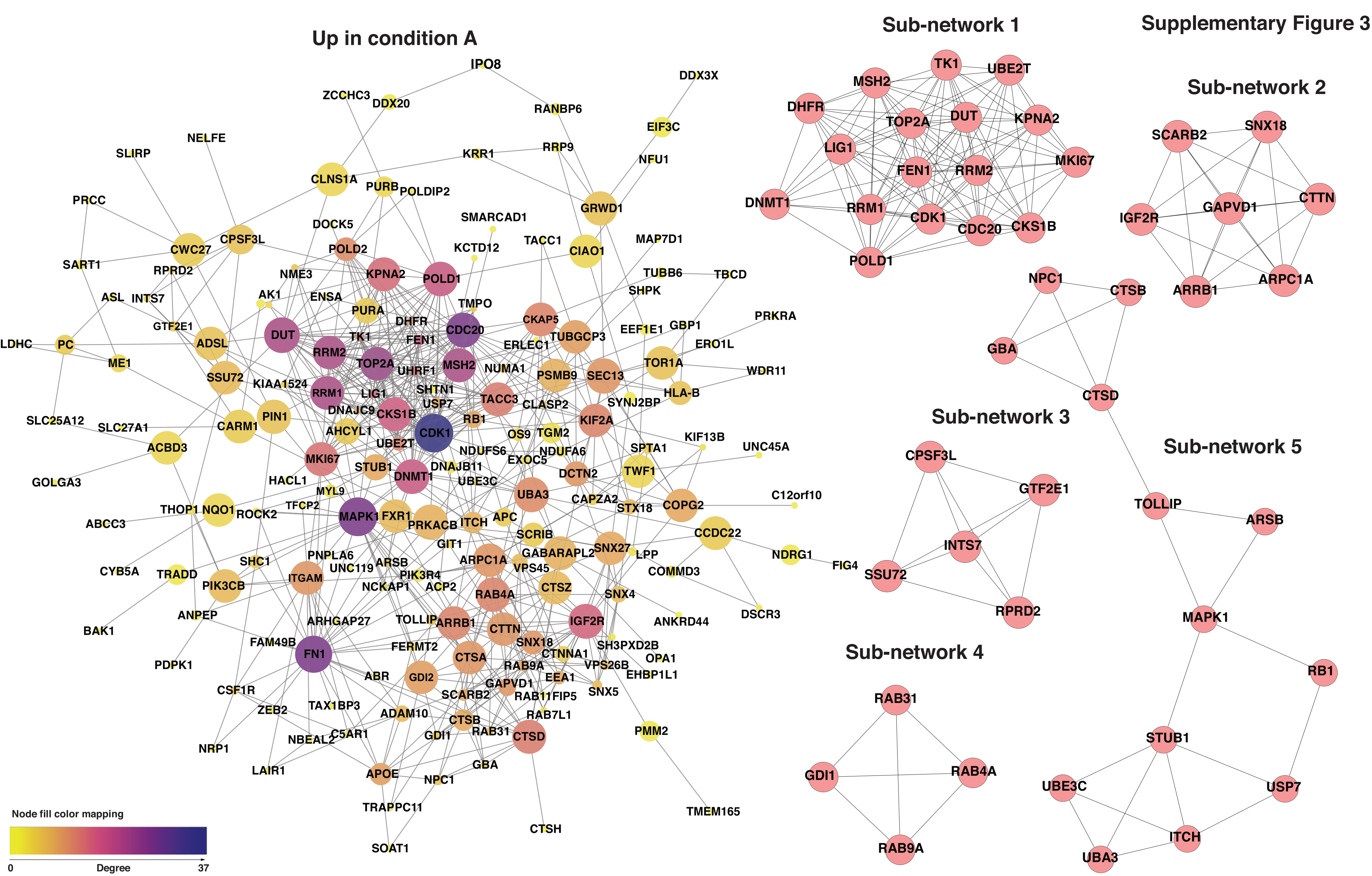


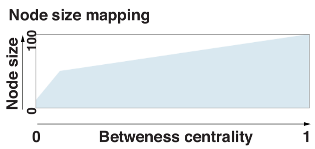


**Supplementary Figure 3**
Network analysis of proteins upregulated in condition A. The network properties were calculated, and the betweenness centrality and degree measures have been indicated using node size and color, respectively. Subclustering of network yielded functional clusters and have been depicted.
