## Supplementary Figure 6 for "Dose-dependent phorbol 12-myristate-13-acetate-mediated monocyte-to-macrophage differentiation induces unique proteomic signatures in THP-1 cells"

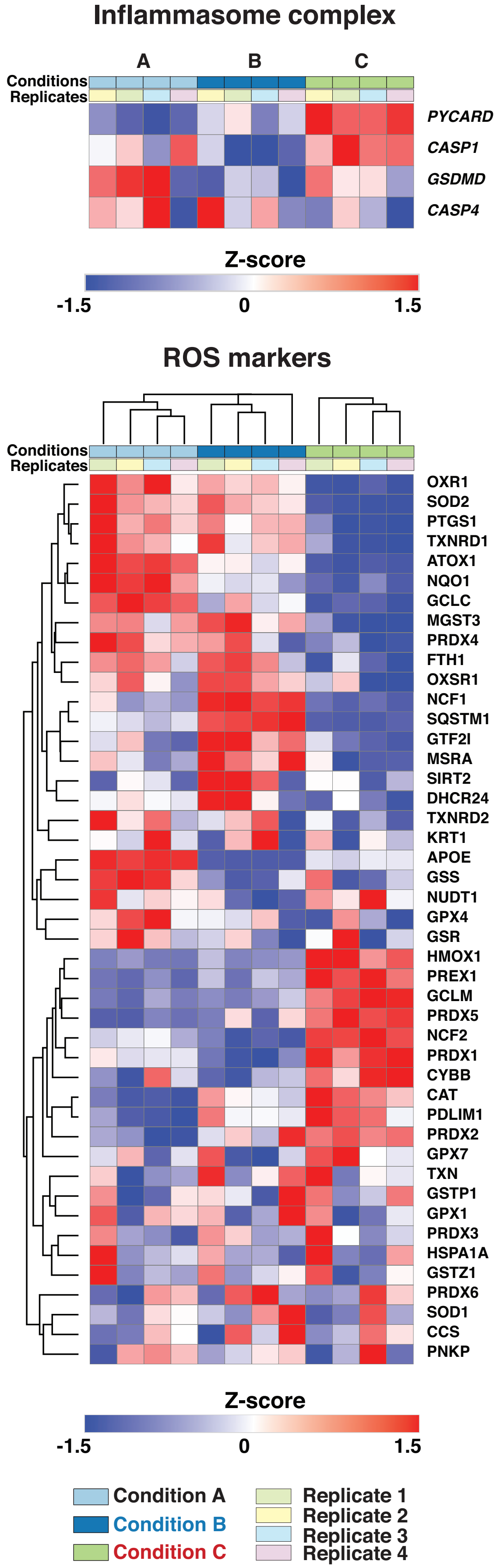


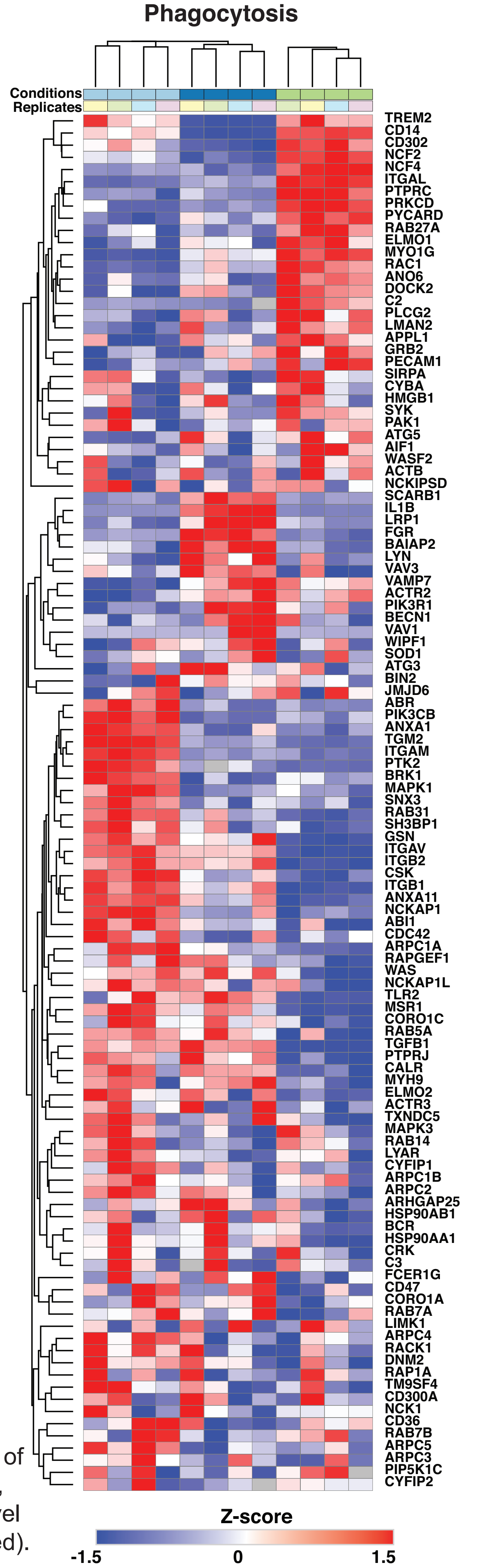


**Supplementary Figure 6**
Dynamic expression pattern to assess the expression levels of proteins that are components of the inflammasome complex, ROS markers, and phagocytosis. The scale indicates the level of expression (Log2-expression values, z-transformed, scaled).
